# Done in 65 ms: Express visuomotor responses in upper limb muscles in Rhesus Macaques

**DOI:** 10.1101/2023.03.02.530807

**Authors:** Aaron L. Cecala, Rebecca A. Kozak, J. Andrew Pruszynski, Brian D. Corneil

**Affiliations:** Department of Physiology and Pharmacology, Western University, London, ON, Canada, N6A 5B7; Robarts Research Institute, 1151 Richmond St. N, London, ON, Canada, N6A 5B7; Graduate Program in Neuroscience, Western University, London, ON, Canada, N6A 5B7; Department of Psychology, Western University, London, ON, Canada, N6A 5B7

**Author notes:** Corresponding Author: Brian D. Corneil: Robarts Research Institute, 1151 Richmond Street North, Western University, London, ON, Canada, N6A 5B7.

**Keywords:** rhesus monkey, visually-guided reaching, electromyography, sensorimotor transformation

## Abstract

How rapidly can the brain transform vision into action? Work in humans has established that the transformation for visually-guided reaching can be remarkably rapid, with the first phase of upper limb muscle recruitment, the *express visuomotor response*, beginning within less than 100 ms of visual target presentation. Such short-latency responses limit the opportunities for extensive cortical processing, leading to the hypothesis that they are generated via the subcortical tectoreticulospinal pathway. Here, we examine if non-human primates (NHPs) exhibit express visuomotor responses. Two male macaques made visually-guided reaches in a behavioral paradigm known to elicit express visuomotor responses in humans, while we acquired intramuscular recordings from the deltoid muscle. Across several variants of this paradigm, express visuomotor responses began within 65 ms (range 48–91 ms) of target presentation. Although the timing of the express visuomotor response did not co-vary with reaction time, larger express visuomotor responses tended to precede shorter latency reaches. Finally, the magnitude of the express visuomotor response was muted on trials where NHPs withheld a reach to one stimulus in order to move to a stimulus appearing 34 ms later in the opposite direction. Overall, the response properties and contextual control of express visuomotor responses in NHPs resemble those in humans. Our results establish a new benchmark for visuomotor transformations underlying visually-guided reaches, setting the stage for experiments that can directly compare the role of cortical and subcortical areas in reaching when time is of the essence.

**Significance statement:** Express visuomotor responses in upper limb muscles are brief periods of recruitment preceding visually-guided reaches. Such responses begin ∼90 ms after visual target presentation in humans, and potentially arise from signaling along the tecto-reticulo-spinal pathway. Here, we show that express visuomotor responses in macaques upper limb muscles resemble those in humans, excepting that they evolve ∼65 ms after target onset, consistent with shorter responses latencies in macaques versus humans. Our results clock the completion of the visuomotor transformation for rapid reaching, and set the stage for experiments to directly test the underlying substrates.

## Introduction

When required, the brain has a remarkable capacity to rapidly transform sensory input into motor output. Expedited responses can consist of eye movements to evaluate a potential threat, or mid-flight adjustments when reaching to a slipping cell phone. In some cases, the brain relies on subcortical circuits that provide a short-latency link between sensation and action, which are nested within broader fronto-parietal circuits mediating more deliberative actions. This appears to be the case for express saccades, which rely on the integrity of the midbrain superior colliculus (SC) and are potentiated by, but not critically reliant on, input from fronto-parietal circuits (Schiller et al., 1987; Edelman and Keller, 1996; Dorris et al., 1997; Dash et al., 2018, 2020). Although the SC has been implicated in mid-flight reach corrections (Day and Brown, 2001; Pruszynski et al., 2016; Reschechtko and Pruszynski, 2020; Suzuki et al., 2022) and interceptions of rapidly moving targets (Perfiliev et al., 2010), debates remain about the comparative role of subcortical versus cortical areas in expedited visually-guided reaching (Fautrelle et al., 2010; Gaveau et al., 2014; Novembre and Iannetti, 2021).

To better understand the neural mechanisms underlying expedited reaches, we and others have studied the timing and response properties of the first change of upper limb muscle recruitment elicited by visual stimulus onset. Doing so establishes the benchmark for how rapidly premotor circuits transform vision into action, and inferences can be drawn by comparing the timing and magnitude of muscle activity to response properties in candidate premotor areas. Such work has revealed that the first wave of upper limb muscle recruitment driven by stimulus presentation evolves within less than 100 ms in humans, regardless of the ensuing reaction time (Pruszynski et al., 2010; Wood et al., 2015; Gu et al., 2016; Kozak et al., 2019; Contemori et al., 2021a). This phenomenon has been termed the stimulus-locked response (Pruszynski et al., 2010) or fast or rapid visuomotor response (Glover and Baker, 2019; Kozak and Corneil, 2021), but lately we have adopted the term express visuomotor response (Contemori et al., 2021a, 2021b) to emphasize similarities with both express saccades and the timing and response properties of visual bursts on premotor cells in the motor layers of the SC (Corneil and Munoz, 2014). The timing and magnitude of such SC visual responses, express saccades, and express visuomotor responses on upper limb muscles are all potentiated by high-contrast, low spatial frequency stimuli (Marino et al., 2012, 2015; Wood et al., 2015; Chen et al., 2018; Kozak et al., 2019; Kozak and Corneil, 2021), and by a variety of cognitive processes promoting premotor preparation (Rohrer and Sparks, 1993; Paré and Munoz, 1996; Munoz et al., 2000; Edelman et al., 2007; Boehnke and Munoz, 2008; Li and Basso, 2008; Bibi and Edelman, 2009; Gu et al., 2018; Contemori et al., 2021b, 2022a).

Here we aim to establish whether express visuomotor responses can be observed in non-human primates (NHPs). We examine three response properties. First, express saccades and the related phenomenon of express neck muscle responses occur ∼20 ms earlier in NHPs than humans (Fischer and Boch, 1983; Fischer and Ramsperger, 1984; Corneil et al., 2004; Goonetilleke et al., 2015). If evoked by a similar mechanism, express visuomotor responses in arm muscles in NHPs should evolve ∼60-80 ms after stimulus onset. Second, although more time-locked to stimulus rather than movement onset, larger express visuomotor responses in humans consistently precede shorter reaction times (Pruszynski et al., 2010; Gu et al., 2016; Contemori et al., 2021a); we predict a similar inverse relationship between recruitment magnitude and reaction time in NHPs. Third, the magnitude of express visuomotor responses in humans is dampened if subjects know to move away from or occasionally not respond to the stimulus (Wood et al., 2015; Gu et al., 2016; Atsma et al., 2018; Kozak et al., 2020; Contemori et al., 2021b, 2022a). Such dampening is a signature of top-down cortical control of a subcortical circuit, and we predict that similar top-down control can be instantiated in NHPs. Overall, we find that NHPs readily generate express visuomotor responses consistent with all these properties, establishing the NHP as a suitable animal model to study underlying neural mechanisms.

## Methods

### Subjects and Surgical Procedures

Two male rhesus monkeys (*Macaca mulatta;* monkey G and monkey B), weighing 8.5 and 13.1 kg respectively, were used in these experiments. All training, surgical, and experimental procedures were in accordance with the Canadian Council on Animal Care policy on the use of laboratory animals (Olfert et al., 1993) and approved by the Western University Animal Use Subcommittee. We monitored the monkeys’ weights daily and their health was under the close supervision of the university veterinarians.

Each animal underwent an aseptic procedure in which a head post was attached to the skull by way of a dental acrylic pedestal prior to behavioral training. During training and experimental sessions, animals were seated with the head-restrained in a customized primate chair that restricted movements of the left arm, torso and hips while permitting unrestricted movements of the right arm in a three-dimensional workspace. Testing was performed in a dimly illuminated room and the monkeys were monitored via a video camera. Visual stimuli, behavioral control, and data acquisition were controlled by MonkeyLogic (Hwang et al., 2019). All visual stimuli were presented on a 42-in. color touch-sensitive monitor (4202L Elo Touch Solutions, Inc., Milpitas, CA) positioned 33 cm in front of the monkeys. The monitor had a spatial resolution of 1920 × 1080 pixels and a refresh rate of 60 Hz.

### The Emerging Target Task

Monkeys generated visually-guided reaches to stimuli presented within the context of an emerging target task (Kozak et al., 2020). The task involves a moving stimulus that temporarily disappears behind an occluder, and then emerges in motion at either the left or right outlet. This task was chosen as it engenders express visuomotor responses in most human subjects (Contemori et al., 2021a, 2021b; Kozak and Corneil, 2021; Kearsley et al., 2022), likely due to the combination of salient visual stimulus presentation following a fixed period of implied motion.

Data was collected from three variants of the emerging target task (**Fig. 1**). All trials started with a red or green ‘start position’ stimulus that appeared at a central location below a gray rectangular occluder. The monkeys were required to touch and hold the location of this start position stimulus within a computer-defined window of 7 cm. Following this, a red or green ‘target’ stimulus that matched the color of the start position stimulus appeared above the occluder. Providing the monkey kept in contact with the start position stimulus for 500 ms, this target stimulus dropped vertically at a constant velocity of 20 cm/s, and then disappeared behind the occluder. After a fixed latency of 300 ms, a red target then emerged in motion below the occluder at either the right or left outlet. At the same time, the start position stimulus disappeared and a secondary visual stimulus that lay under a photodiode (and hence was unseen by the monkey) was presented for data alignment purposes. On double target trials, a green target emerged in motion at the other outlet 34 ms later. To obtain a reward, the monkeys had to maintain contact on the start position until target emergence below the barrier, and then reach to touch the colored target that matched the color of the stimuli presented at the start of the trial before it disappeared off the screen (i.e., reach to the red target on single target red (**Fig. 1A**) and double target red (**Fig. 1B**) trials, or to the green target on double target green trials **(Fig. 1C)**). Thus, correct performance on double target green trials required the monkeys to not reach towards the red target that emerged slightly (34 ms) earlier. The moving target that appeared below the barrier started 15 cm to the right or left of the start position stimulus, and then moved in an lateral-downward direction (45 deg below horizontal) at 20 cm/s. Trials where the monkey touched or moved toward the wrongly colored target were tabulated as incorrect reaches, and retained for further analysis. We decided to emphasize response speed over accuracy, and hence used large acceptance windows (radius of 11 cm) around the emerging target. An analysis of endpoints showed that both monkeys tended to reach more accurately than this window, with the standard deviations of horizontal or vertical endpoints being 2.5 and 1.9 cm for monkey G and 1.5 and 1.0 cm for monkey B. Recent work in humans (Contemori et al., 2022b) shows that larger express visuomotor responses are observed when subjects land accurately on target, as opposed to deliberately overshooting or undershooting the target. Thus, we do not believe our results are influenced by our choice to emphasize response speed over accuracy.

**Figure 1.**
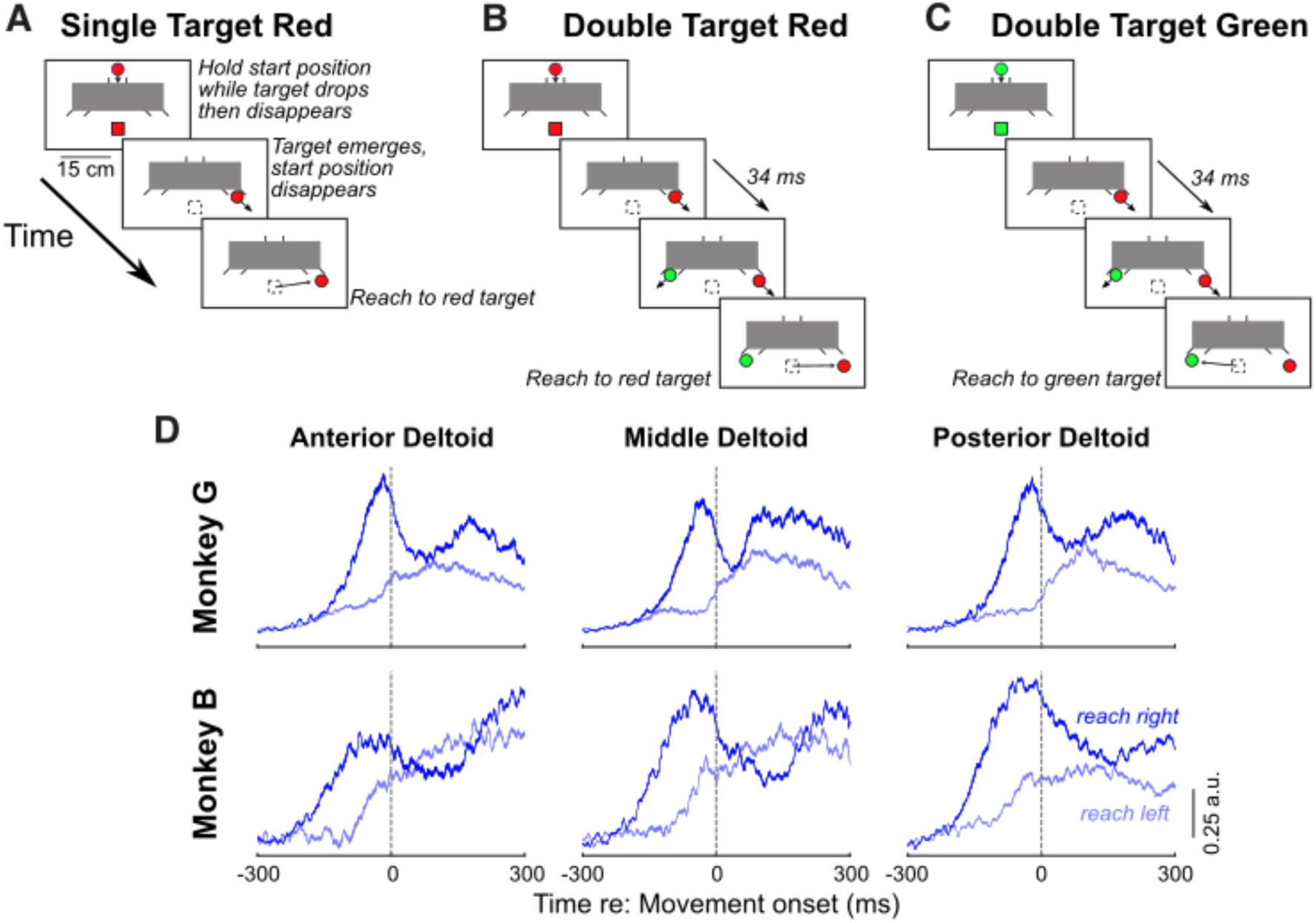
**A-C.** Emerging Target Task. At the start of each trial, either a red (**A, B**) or green (**C**) stimulus appears at the start position (red or green squares), below an occluder (gray box). The monkey touches and holds the screen at this start position. A target of the same color then appears above the occluder and moves down the chute before disappearing briefly behind the occluder for 300 ms. In Single Target Red trials (**A**), a single red target emerges in motion below the occluder at one of two outlets. In the Double Target Red (**B**) and Double Target Green (**C**) trials, appearance of the red target in one outlet was followed 34 ms later by appearance of a green target at the other outlet. The monkey was rewarded for reaching to the target that matched the color of the stimuli presented at the start of the trial. **D.** Movement-aligned activity for the anterior, middle, and posterior heads of deltoids for the two monkeys for single-target red trials, showing greater recruitment for rightward than leftward reaches. Data shows the average of the mean recruitment from each session, normalized to the peak movement-aligned activity from each session. Contours subtend the mean ± stderr.

Within a given block of trials, all trial conditions (defined by target location and, when used, task variant) were presented with equal frequency but randomly shuffled. Some datasets consisted of only single-target red trials (2 sessions for monkey G, 1 session for monkey B). For the other days, collected data in a block of ∼100 single-target red trials, prioritizing such data as EMG recording electrodes occasionally were removed as animals performed the task (this happened in one day for monkey G). After these 100 single-target red trials, we then intermixed the other task variants. Across all experimental sessions, we obtained muscle recordings from a minimum of 380 to a maximum of 1470 trials.

### EMG electrode insertion and data collection

Prior to a data collection session, monkeys were lightly sedated before intramuscular, fine-wire EMG electrodes were acutely inserted into the posterior, medial, and anterior heads of the deltoid muscle. Sedation followed an established procedure (Corneil et al., 2012), involving induction with a low-dose of medetomidine, and reversal after electrode insertion with atipamazole. After reversal, monkeys were given ∼20 minutes to recover prior to beginning data collection. For each intramuscular recording, we inserted two electrodes into the muscle belly staggered by ∼1 cm to enable bipolar recordings of bulk muscle activity. Each electrode was fabricated in-house by threading PFA-coated 7-strand stainless steel wire (Type 316, A-M Systems, Sequim, WA) into a 27 gauge cannula, and then stripping the final ∼3 mm of insulation. Anatomically, the different heads of the deltoid were identified by manually palpating the head of the humerus. A ground electrode was positioned on the dorsal neck approximately between the scapulae. Intramuscular EMG activity was differentially amplified and recorded with a Myopac Junior system (Run Technologies, low-pass filter modified to 2 kHz) and digitized at a rate of 1,000 Hz onto a Plexon Omniplex system, which also received the photodiode signal. Off-line, any DC offset was removed and EMG data were full-wave rectified and smoothed with a 7-point running average. Our sampling frequency was limited to 1,000 Hz, which undersamples intramuscular signals. We confirmed that such undersampling had only a minor influence on our results by collecting data from one additional session on a Ripple Grapevine system that sampled EMG data at 30,000 Hz. This allowed us to compare results obtained from 3 simultaneously recorded muscles processed offline by either downsampling the EMG signal to 1,000 Hz versus that obtained by first full-wave rectifying the 30 kHz signal followed by integration into 1 ms bins (Loeb and Gans, 1986; Elsley et al., 2007). For all three muscles recorded at 30 kHz, these different processing steps changed the discrimination latency of the express visuomotor response by only 1 ms. Thus, while our undersampled data would not be appropriate for identifying single motor-unit action potentials, it does accurately capture the timing of changes in multi-unit recruitment after visual target presentation.

### Data Analysis

All data analyses were conducted in MATLAB^®^ (R2021a; Natick, MA, USA). We rejected trials where the monkey either did not respond or failed to respond within 500 ms, as well as trials where movements began within less than 80 ms of target emergence below the barrier. Reach RT was calculated as the time from target emergence below the barrier (indicated by the photo-diode signal) to the time point where the monkey’s hand was either no longer in contact with the touchscreen at the start position, or when the monkey’s hand started deviating toward the emerging target. Offline, we further inspected all trials within a graphical user interface, allowing us to adjust the movement onset mark at a resolution of 1 kHz, and flag trials where the initial movement was in the opposite direction of the correct response. Monkey G moved more rapidly and tended to remain in contact with the screen during the reach (i.e., ‘smearing’ their hand across the screen), whereas Monkey B moved more slowly and had a greater tendency to lift their finger off the screen when reaching. Thus, incorrect reaches in Monkey G included trials where the monkey first moved in the incorrect direction before correcting their reach in mid-flight. In contrast, incorrect reaches in Monkey B were sometimes only identified when the monkey touched the wrong location. On other occasions, an incorrect reach in Monkey B was identified by a deflection in the horizontal touch screen position in the wrong direction, as identified in the user interface.

The different heads of the deltoid muscle are known to have different preferred reaching directions in the rhesus macaque (Kurtzer et al., 2006). Note however that the reaching task in this study required that the monkeys reach forward and maintain contact with the touchscreen at the start point, and then upon target emergence move their arm to the right or left to obtain a reward. In the context of this bidirectional task, as shown in **Fig. 1D**, all heads of deltoid exhibited greater recruitment before rightward versus leftward reaches on the touchscreen. Therefore, for the purpose of analyzing the express visuomotor response, the rightward direction was deemed to be the preferred direction of all muscles.

As with previous work (Pruszynski et al., 2010; Gu et al., 2016; Contemori et al., 2021a), we used a time-series receiver-operating characteristic (ROC) analysis to define the presence and latency of the express visuomotor response. This analysis method is suitable when targets can appear at one of two diametrically opposite locations, although we note that express visuomotor responses have been studied with other methods with different potential target configurations (Gu et al., 2019; Kearsley et al., 2022; Selen et al., 2023), and with single-trial analyses (Contemori et al., 2021b, 2022a). Data recorded from each muscle was treated as an independent sample, and data was segregated by side of target emergence and task variant. For blocks with multiple interleaved task variants, separate time-series ROCs were run on data for each task variant, comparing EMG activity after left or right target emergence. The ROC value (i.e., the area under the ROC curve) was calculated and plotted at every time sample (1 ms) from 100 ms before to 500 ms after target emergence (e.g., **Fig. 2C**); this value indicates the likelihood of discriminating the side of target emergence based on EMG activity alone. A ROC value of 0.5 indicates chance discrimination, whereas values of 1 or 0 indicate perfectly correct or incorrect discrimination, respectively. Although time-series ROCs often fluctuated around 0.5 in the 100 ms surrounding target emergence, we often noted the presence of linear trends in this interval as well as a high degree of variance on task variants with fewer overall trials. Such trends have been observed previously in previous studies (Goonetilleke et al., 2015), and we suspect they arise from trials with short reaction times where the animal was guessing, which are included or excluded if the animal guessed correctly, or not. These linear trends precluded the use of fixed thresholds to determine the latency of any express response. Accordingly, we first detrended the time-series ROC based on a linear regression run on the time-series ROC over a baseline interval spanning from 100 ms before to 45 ms after target emergence. We then set a discrimination threshold to be the first of 8 of 10 points where the time-series ROC exceeded 3 standard deviations above the mean of the detrended baseline time-series ROC. If this time was between 45 to 100 ms after target emergence, then the muscle was deemed to be exhibiting an express visuomotor response, and the latency was taken to be the discrimination time. The magnitude of the express visuomotor response was determined by the integral of EMG activity measured over the next 30 ms. All EMG data are reported in source microvolts, and were not subjected to any normalization unless otherwise noted.

**Figure 2.**
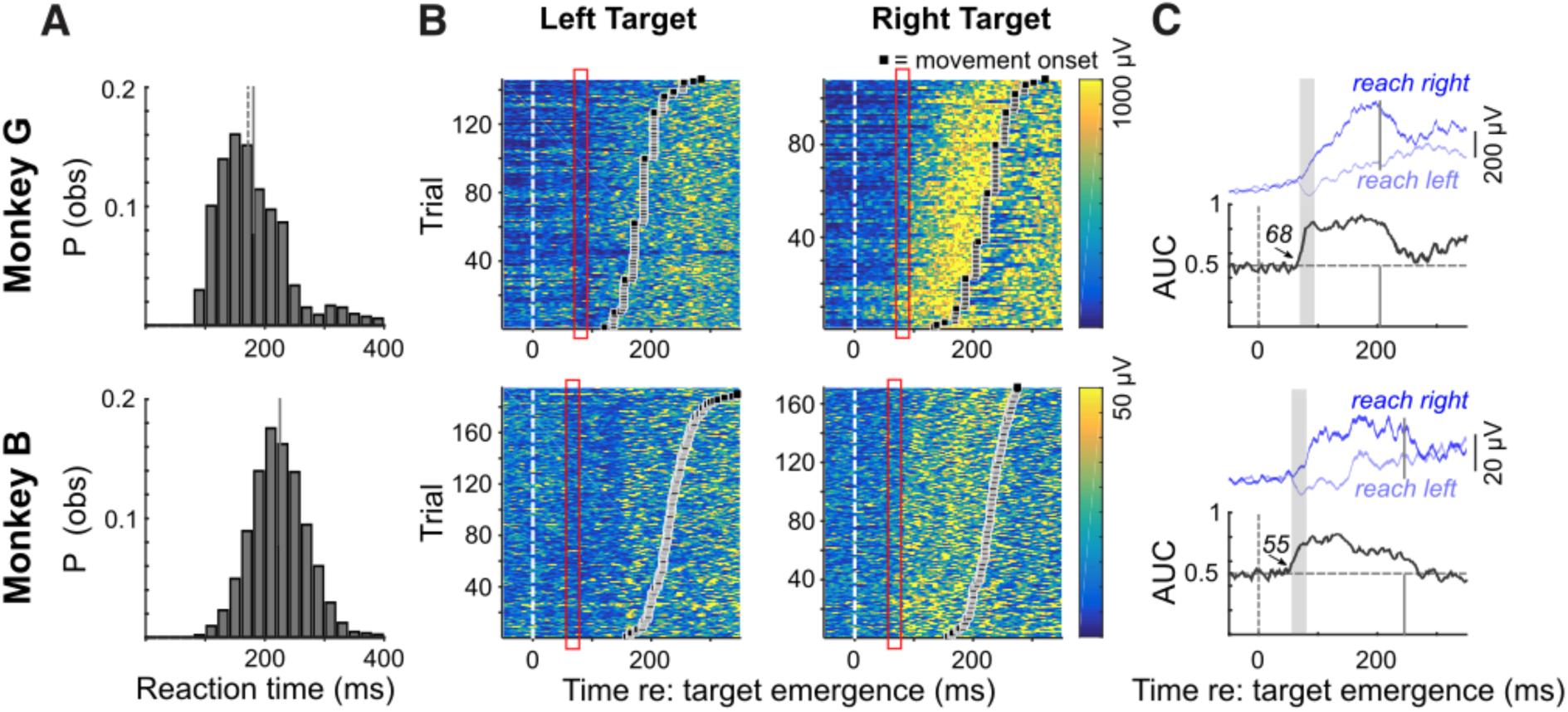
**A.** Frequency histograms of reaction times (20 ms bins) for Monkey G (top row, n = 1303 trials) and Monkey B (bottom row, n = 2648 trials) across all experimental sessions. Solid/dashed lines depict means/medians. **B.** Representative examples of express visuomotor responses in Monkey G (top row) and Monkey B (bottom row), depicting trial-by-trial EMG recruitment aligned to target emergence to the left or right. In these plots, trials are sorted by movement onset (black squares), and the color conveys the magnitude of recruitment. **C**. For each monkey, the top row depicts mean EMG activity subtended by the standard error for left (light blue) and right (dark blue) reaches, and bottom row shows the time-series ROC (AUC = area under curve). Solid vertical lines depict mean reaction time for the representative examples. The red rectangles in **B** and gray rectangles in **C** show the 25 ms interval following the detection of the express visuomotor response.

### Experimental design and statistical analyses

All statistical analyses were performed in MATLAB^®^ (R2021a; Natick, MA, USA). Comparisons of the magnitude and latencies of express visuomotor responses and of reaction times across the various trial types were performed via *t*-tests, using paired and unpaired *t*-tests when appropriate. Correlational analyses were performed on latency, magnitude, and RT measures, as described below. Comparisons of error rates across different task variants were performed using a chi-squared test.

## Results

Two rhesus macaques reached to red or green targets presented in the context of the emerging target task (**Fig. 1A-C**). This task readily elicits express visuomotor responses in humans (Kozak et al., 2020; Contemori et al., 2021a; Kozak and Corneil, 2021). Our goal was to establish the phenomenon of express visuomotor responses in monkey, by: i) describing the response latencies and consistency of such responses both within and across experimental sessions, ii) investigating the relationships with ensuing reach reaction times (RTs), and iii) establishing whether the express visuomotor response to the red stimulus would be decreased when monkeys were instructed to reach in the opposite direction to a delayed emerging green stimulus. Our dataset consists of EMG recordings made in 8 sessions in monkey G (7, 8, and 7 successful recordings from anterior, medial, and posterior deltoid respectively) and 10 sessions in monkey B (9, 8, and 8 from anterior, medial, and posterior deltoid respectively). As all muscles exhibited greater recruitment for rightward vs leftward reaches on the touchscreen (**Fig. 1D**), and since we noted no differences in the express visuomotor responses from the three heads of deltoid, we treated each recording as an independent sample of multiple motor units, and pooled data within a given monkey.

### Stimulus emergence evokes short-latency express visuomotor responses

We begin by describing the behavior and patterns of muscle recruitment immediately following target emergence in single target red trials. On such trials, Monkey G reached within 179 ± 61 ms (median = 170 ms) and Monkey B reached within 223 ± 47 ms (median = 220 ms) (**Fig. 2A**). Both animals made some reaching errors when only a single red target emerged (6% for Monkey G, 5% for Monkey B), but more commonly anticipated target emergence (8.6% for Monkey G, 12.1% for Monkey B). Representative examples of EMG data recorded from each monkey are shown in **Fig. 2B** and **C**. As animals held their arm against gravity in order to touch the central starting position, there was a modest level of recruitment prior to and immediately after target emergence. Very soon after target emergence, muscle recruitment increased or decreased following target emergence to the right or left, respectively. Such lateralization is a hallmark of express visuomotor responses in humans (Pruszynski et al., 2010; Wood et al., 2015), and emphasizes that express visuomotor responses are tuned to stimulus location rather than being a generic startle or preparatory response (Gu et al., 2019; Selen et al., 2023). Another defining feature of express visuomotor responses in humans is the trial-by-trial consistency in which this initial change in muscle recruitment is more locked to target emergence rather than movement onset. This indeed was the case in the representative examples shown in **Fig. 2B** (see red rectangles): note how this initial lateralized change in muscle activity occurred ∼60-70 ms after target emergence, well before the hand moved or lifted away from the starting location. Following the express visuomotor response, rightward reaches were preceded by a gradually increasing level of recruitment that peaked just before reach initiation, whereas muscle recruitment before leftward reaches persisted near or below the baseline level of activity prior to target emergence. The data shown for Monkey B exhibited a second target-locked decrease in muscle activity ∼130-140 ms after left target emergence, resembling reports of oscillations in muscle recruitment initiated by a visual stimulus in humans (Wood et al., 2015). Such double-dips were seen in about half of the recordings in Monkey B but were not observed in Monkey G. We suspect this is because monkey G had generally shorter RTs, so that the recruitment aligned with reach onset overlapped with any subsequent activity following the express visuomotor response.

To analyze the timing of express visuomotor responses, we performed a time-series receiver operating characteristics (ROC) analysis. This analysis determines the timing at which information following leftward or rightward target emergence influenced muscle recruitment. For the examples shown in **Fig. 2C**, the express visuomotor response began 68 ms (Monkey G) or 55 ms (Monkey B) after target emergence. In both examples, the time-series ROC rose quickly after this discrimination time, and remained elevated for the next ∼200 ms.

We repeated this analysis across our sample of deltoid recordings. Across our sample, an express visuomotor response, which was defined as occurring within 45-100 ms, was detected in 77% (17 of 22; 7, 8, and 7 recordings from anterior, middle, and posterior deltoid) recordings in monkey G and in all 25 recordings in monkey B (9, 8, and 8 recordings from anterior, middle, and posterior deltoid). When present, express visuomotor responses began very soon after stimulus emergence, with discrimination latencies being 65 ± 8 ms (range: 52 to 81 ms) for monkey G and 64 ± 11 ms (range: 48 to 91 ms for monkey B; **Fig. 3A**; no difference between monkeys, *t*-test, *t*(40) = 0.34, *p* = 0.72). A potential concern about these short discrimination latencies is that they may arise only because of recruitment associated with short-RT reaches. We therefore repeated the time-series ROC analysis across our sample only on the slower half or all trials. Excluding the shorter-than-average RT trials in each session increased reach reaction times by ∼25 ms to 205 ± 63 ms for Monkey G and 253 ± 37 ms for Monkey B. Express visuomotor responses persisted in 72% (16 of 22) of recordings in monkey G and in all 25 recordings in monkey B, with average discrimination latencies being at 69 ± 7 ms for monkey G and 68 ± 12 ms for monkey B (**Fig. 3A**). Although significant in both monkeys (paired *t*-test, monkey G: *t*-test, *t*(15) = −6.9, *p* < 10^−5^; monkey B: *t*(24) = −3.10, *p* = 0.005), the ∼4 ms increase in discrimination latency after excluding shorter-than-average RTs was less than the ∼25 ms increase in mean RT. We suspect that slightly longer discrimination latencies reflects lower magnitude express visuomotor responses associated with longer RT trials (see next section), as reported previously (Contemori et al., 2021a, 2021b).

**Figure 3.**
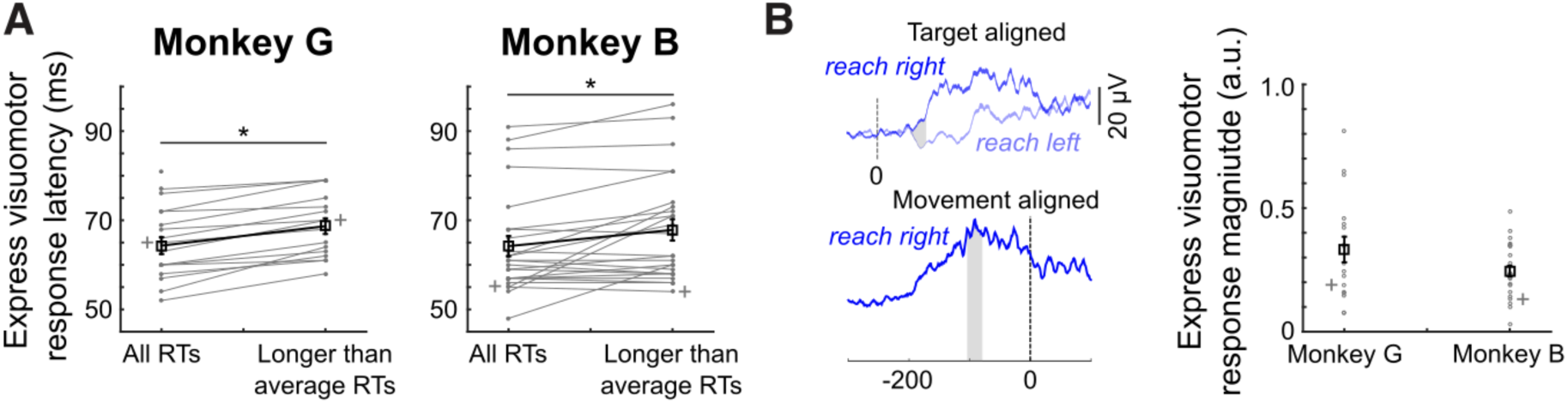
Properties of express visuomotor responses. **A.** Latencies of express visuomotor responses for both monkeys, showing latencies when all trials are considered regardless of reaction time, and after including only the longer-than-average reaction times. Gray lines connect observations from the same muscle from a given experimental day (grey + marks denote representative data shown in Fig. 2). Black lines and squares show mean ± standard error. Asterisks show significant differences of paired t-test (p < 0.05). **B**. Depiction of the calculation of the proportional magnitude of the express visuomotor response, using the representative data from Monkey B. Using data normalized to the peak of movement-aligned activity, we integrate the difference in the mean EMG curve for right versus left responses over the 25 ms after the onset of the express visuomotor responses, and divide this by the integral of the 25 ms interval surrounding peak EMG activity aligned to right reach onset. Same general format as **A**; asterisk showing significant results of a student’s t-test (p < 0.05).

Previous work in humans has shown that express visuomotor responses observed in the emerging target task can be quite large in magnitude, occasionally equalling if not exceeded ‘voluntary’ recruitment magnitude achieved when recruitment is aligned to reach onset (Contemori et al., 2021a; Kozak and Corneil, 2021). To establish the proportional magnitude of the express visuomotor response, we expressed it relative to a 25 ms interval surrounding the peak magnitude of movement-aligned recruitment (**Fig. 3B**). Expressed in this fashion, the express visuomotor response reached 0.33 ± 0.21 (range: 0.08 to 0.81) for monkey G and 0.25 ± 0.11 (range: 0.03 to 0.48) for monkey B (proportional magnitudes were not significantly larger in monkey G than monkey B; *t*-test, *t*(40) = 1.76, *p* = 0.08). Thus, on average, the recruitment magnitude of the express visuomotor response was about one quarter to one third of the magnitude of reach-related recruitment.

### Express visuomotor response magnitude inversely correlates with RT

Although the express visuomotor response may be modest in magnitude, previous work shows that larger express visuomotor responses tend to precede short RTs (Pruszynski et al., 2010; Gu et al., 2016; Contemori et al., 2021a). To investigate whether a similar inverse correlation is seen in NHPs, we calculated the trial-by-trial magnitude of EMG recruitment during an express visuomotor response window, integrated over 25 ms after the express visuomotor response was detected (Pruszynski et al., 2010), and plotted it as a function of RT. **Figure 4** shows these plots for the representative data from monkey G (**Fig. 4A**) and monkey B (**Fig. 4B**), and both show the inverse correlation between magnitude and RT expected from previous studies (Monkey G: *r* = −0.56, *p* < 10^−9^; Monkey B: *r* = −0.37, *p* < 10^−6^). We repeated this analysis for every session in our sample, and for both monkeys, observed the expected negative correlation (**Fig. 4C**; Monkey G: mean *r =* −0.45 ± 0.26. Monkey B: mean *r* = −0.29 ± 0.16). The distributions for both monkeys lay significantly below zero (*t*-test; Monkey G: *t*(16) = −7.3, *p* < 10^−5^. Monkey B: *t*(24) = −9.4, *p* < 10^−8^).

**Figure 4.**
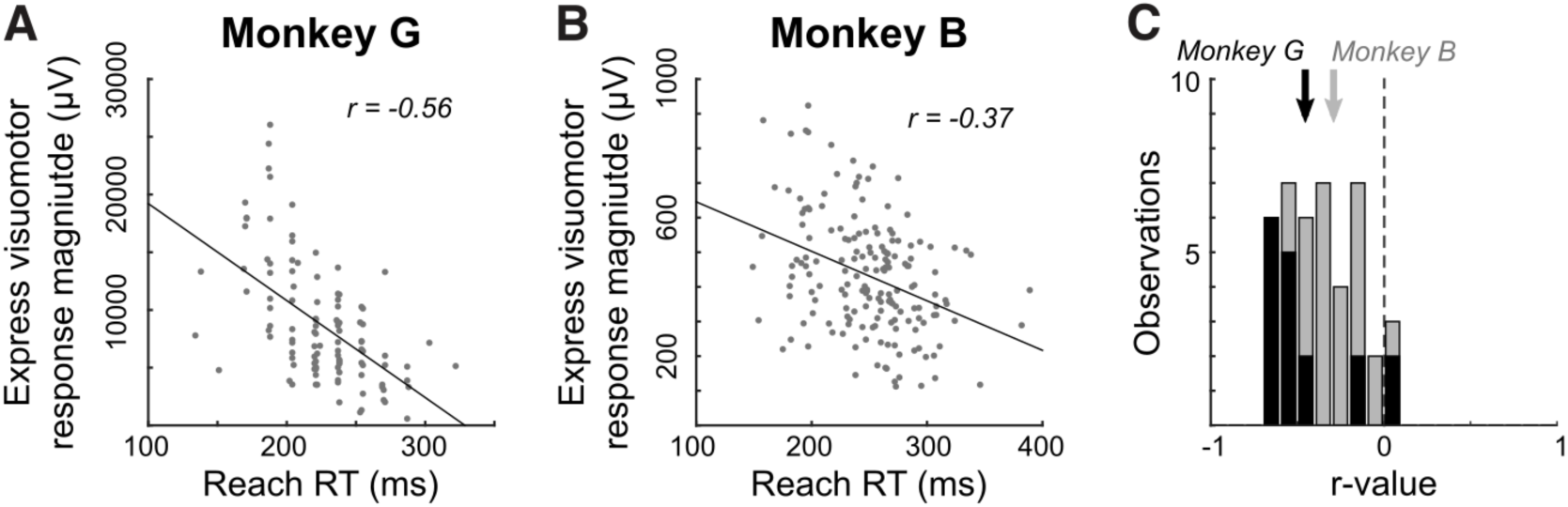
Larger express visuomotor responses tended to precede shorter reach reaction times (RTs). **A, B**. Representative data plotting magnitude of express visuomotor response as a function of reach reaction time, from the representative sessions shown in Figure 2. Each data point represents a single trial, and the black diagonal line shows the linear regression. The correlation coefficient (*r*) is shown in the top-right of each subplot. **C**. Histogram of correlation coefficients across all sessions in which an express visuomotor response was detected. Vertical dash line indicates 0. Solid or gray lines or bins denote data for Monkey G or B, respectively. Vertical arrows show mean r-values.

### Contextual control influences the magnitude but not timing of the express visuomotor response

Having established that macaque upper limb muscle recruitment following the emergence of a single target resembles that observed in humans, we now turn to data recorded when two targets emerged below the barrier. The sequence of target emergence was the exact same on double-target trials; the red stimulus always appeared 300 ms after the initial descending stimulus disappeared behind the barrier, and the green stimulus always appeared 32 ms after the red stimulus. Monkeys reached to the emerging stimulus that matched the color of the central hold stimulus and dropping stimulus (**Fig. 1B,C**). Double target green trials therefore required monkeys to not reach to the red emerging stimulus in order to reach to the green stimulus that subsequently emerged on the opposite side. Thus, monkeys had to exert a different degree of contextual control in how they responded to the red emerging stimulus on double target red than double target green trials.

We first compared reaching behavior on double target red vs double target green trials. RTs for correct responses (timed to the emergence of the red stimulus) were longer for double target green trials than double target red trials (**Fig. 5A**, Monkey G: double target red = 188 ± 70 ms, n = 572; double target green = 209 ± 97 ms, n = 206; *t*(776) = −3.36, *p* < 0.001. Monkey B: double target red = 235 ± 50 ms, n = 1607; double target green = 255 ± 49 ms, n = 962; *t*(2579) = −9.63, *p* < 10^−10^). Both monkeys made reaching errors at a significantly higher rate on double target green than double target red trials (Monkey G: chi square-test, *X*^2^ (1) = 33.9, *p* < 10^−8^; Monkey B: *X*^2^ (1) = 11.25; *p* < 10^−3^), and Monkey G generated more errors than Monkey B on double target green trials (chi-square-test, *X*^2^ (1) = 5.21; *p* = 0.02). When generated, such errors started earlier than correct responses on either double target green or double target red trials (Monkey G: error RTs on double target green = 172 ± 45 ms, n = 207; *t*-test against RTs of correct responses on double target green trials *t*(411) = 5.01, *p* < 10^−7^; *t*-test against RTs of correct responses on double target red trials *t*(777) = 3.06, *p* < 0.005; Monkey B: error RTs on double target green = 186 ± 50 ms, n = 409; *t*-test against RTs of correct responses on double target green trials *t*(1376) = 23.38, *p* < 10^−10^; *t*-test against RTs of correct responses on double target red trials *t*(2019) = 17.62, *p* < 10^−10^).

**Figure 5.**
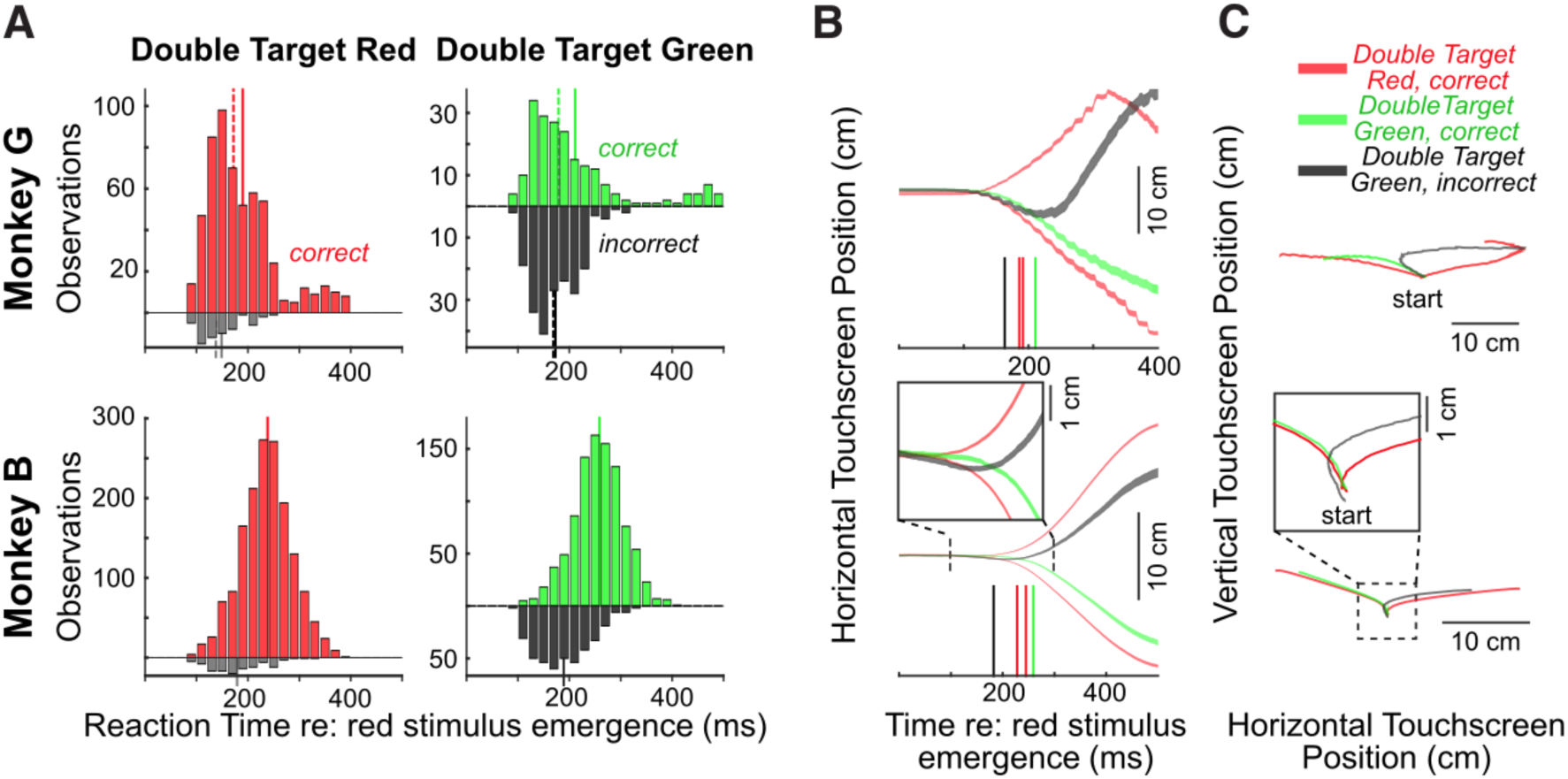
Contextual control on Double Target trials. **A.** Histograms across all sessions of reaction times (20 ms bins) relative to the emergence of the red stimulus for Monkey G (top row) and Monkey B (bottom row), for Double Target Red trials (left column, Monkey G: 632 trials, error rate = 9.4%; Monkey B: 1788 trials, error rate = 9.8%) or Double Target Green trials (right column, Monkey G: 413 trials, error rate = 50.1%; Monkey B: 1378 trials, error rate = 29.6%). Upward/downward histograms represent correct/incorrect responses. Solid/dashed lines depict histogram means/medians. **B.** Trajectory of horizontal touchscreen position averaged across all trials for Double Target Red trials to the right or left (red contours), and Double Target Green trials, showing correct trials when the monkey should have reached left (green contours), and incorrect trials when the monkey should have reached right (black contours). Note how incorrect trials initially go left before being corrected mid-flight to go to the right. Contours show mean ± stderr. Solid colored vertical lines in B depict mean reaction time for corresponding trial type. **C.** Same data, depicting vertical position as a function of horizontal position. Insets in **B** and **C** for Monkey B show zoomed-in representation of early reach trajectory, given the modest movement in the wrong direction.

Both monkeys generated a high proportion of errors on double target trials, leading us to wonder about the degree to which they were exerting contextual control on these trials. First, we did find that the RTs on double target red trials were significantly longer than the RTs on single target red trials, suggesting the presence of intermixed trials slowed the animals down (Monkey G: t(1873)=2.75, *p* < 0.01; Monkey B: t(4258)=8.58, *p* < 10^−10^). Second, while the RTs of both correct and incorrect trials pooled together did not differ significantly on double target green versus double target red trials in either monkey (Monkey G: t(1043)= −1.41, *p* = 0.16; Monkey B: t(3015)= −1.63, *p* = 0.10), incorrect reaches were quite small, and an analysis of the kinematics of reaching revealed notable differences in performance on double target red vs double target green trials. Recall that Monkey G tended to smear his hand on the touchscreen, which allowed us to visualize the trajectory of the reaching response. The grouped trajectories for a subset of reaching responses on double target trials are shown for Monkey G in **Fig. 5B** and **C** (top row). Correct leftward reaches on double target green trials (green traces) follow the trajectory of correct leftward reaches on double target red trials (red traces). In contrast, while incorrect right reaches on double target green trials initially follow a similar trajectory as these leftward reaches (black trace), they are rapidly reversed in direction, and end up at similar locations to correct rightward reaches on double target red trials. Thus, even though Monkey G initiated correct and incorrect movements at similar latencies, later parts of incorrect reaches were rapidly re-directed in the appropriate direction. Similar tendencies are observed in Monkey B, although to a much more subtle degree since Monkey B tended to lift his hand off the touchscreen (see insets zooming in on the early parts of the reach trajectories). These behavioral patterns are consistent with the animals trying, and often failing, to exert more contextual control on double target green trials. However, later parts of the reach trajectory on double target green trials confirmed that both animals had consolidated the task instruction to reach to the correct location.

We next asked the degree to which contextual control influenced the express visuomotor response to the red stimulus. In humans, express visuomotor responses evolve at the same time but are dampened in magnitude when subjects move away from rather than toward a peripheral stimulus (Gu et al., 2016; Kozak et al., 2020), or exert some other degree of contextual control (Wood et al., 2015; Atsma et al., 2018; Contemori et al., 2021b). An analysis of contextual control on the express visuomotor response requires that we compare muscle recruitment on trials where the red stimulus appears at the same location on either double target red or double target green trials. **Figure 6A** shows the mean ± stderr representations of EMG activity from the same representative sessions as shown in **Figure 2**, depicting right middle deltoid recruitment when the red stimulus appeared to right. The monkeys reached to the right on double target red trials (red contours in **Fig. 6A**), and the express visuomotor response resembled that seen on the single target red trials (**Fig. 2B,C**). On double target green trials, the correct response was to withhold the rightward reach, and reach to the left. When the animals did this correctly, the initial express visuomotor response to the red stimulus was muted (green contours in **Fig. 6A**). In contrast, when the monkeys incorrectly reached to the red stimulus on double target green trials, initial muscle recruitment during the express response interval resembled that on double target red trials (compare black and red contours in **Fig. 6A**), whereas later phases of recruitment resembled correctly-performed double target green trials, as the animal corrected their reach. Thus, for these examples, the express visuomotor response to the earlier appearing red stimulus depended both on the context of the trial (double target red vs double target green), and on performance on double target green trials.

**Figure 6.**
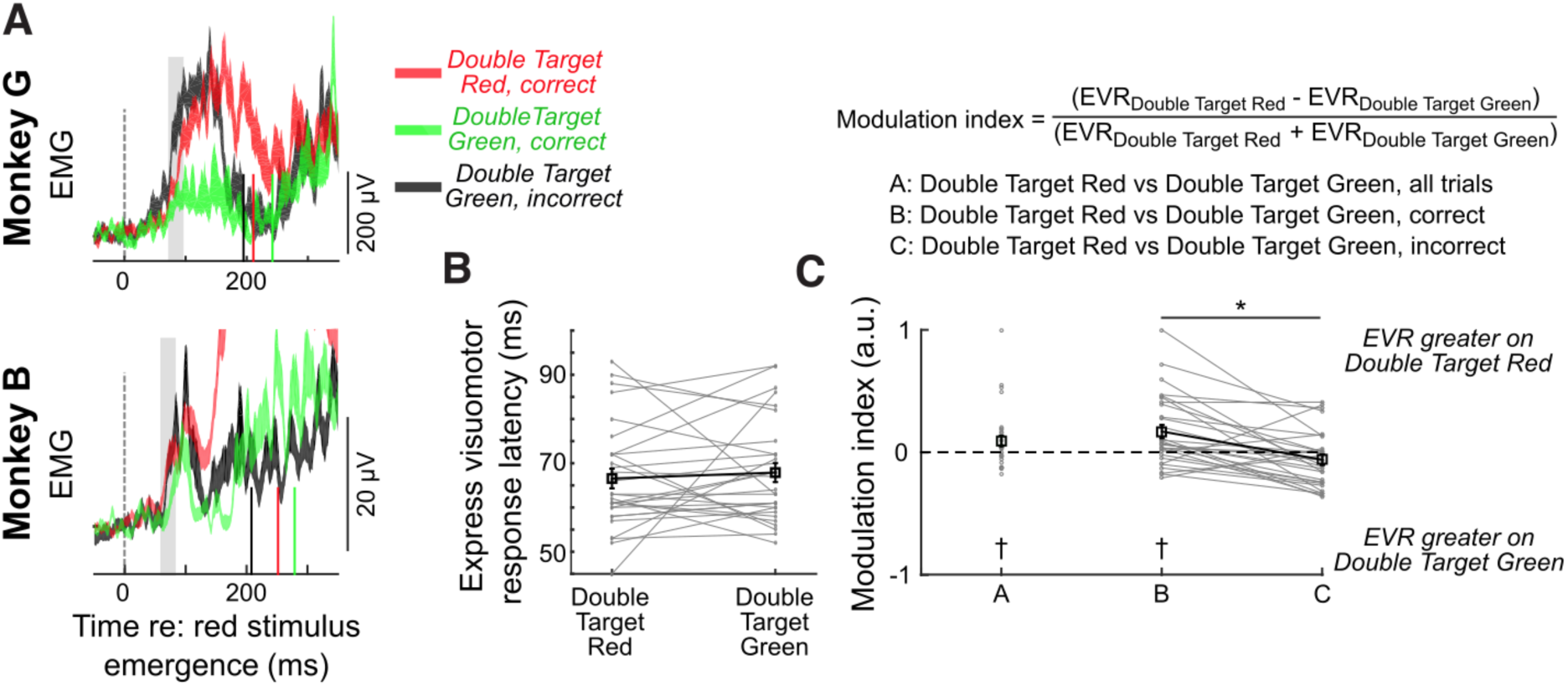
Contextual control on Express Visuomotor Response (EVR) on Double Target trials. **A.** Depiction of mean EMG activity +/− stderr from representative sessions when the red stimulus emerged to the right. Gray rectangles denote the 25 ms interval following the express visuomotor response, over which magnitude was calculated. Solid colored vertical lines in **A** depict mean reaction time for corresponding trial type from representative sessions. **B,C**. The latency (**B**) or magnitude (**C**) of the express visuomotor response, by trial type and/or performance, across 29 occurrences where an express visuomotor response could be detected on both Double Target Red and Double Target Green trials. Magnitudes are expressed as a modulation index (see equation), for a variety of contrasts. Gray lines connect observations from the same muscle from a given experimental day. Black lines and squares show mean +/− standard error. Crosses denote distributions significantly different from zero (t-test, p < 0.05). Asterisks show significant differences of Bonferroni corrected paired t-test (p < 0.05).

Across our sample, we found that contextual control influenced the magnitude but not timing of the express visuomotor response to the red stimulus. To quantify this, we first compared the timing of the express visuomotor response, when such a response was detected on both double-target red and double-target green trials. We found no influence of context on express visuomotor response latency, with the latency being unchanged across double-target red trials and double-target green trials (**Fig. 6B**. Monkey G: double-target red = 66 ± 9 ms; double-target green = 70 ± 14 ms; paired *t*-test; *t*(9) = −0.75, *p* = 0.46; Monkey B: double-target red = 67 ± 13 ms; double-target green = 67 ± 10 ms; *t*(18) = 0.02, *p* = 0.98).

Next, we determined the comparative magnitude of the express visuomotor response over the next 25 ms on double target red or double target green trials, expressing the change in magnitude as a modulation index (see equation in **Fig. 6C**). Calculated this way, a value of 0 arises when there is no modulation of the express visuomotor response magnitude for different trial types, whereas values greater than (less than) 0 arise when the magnitude of the express visuomotor response is larger (smaller) on double target red than double target green trials. Our first calculation (comparison “A” in **Fig. 6C**) compared double target red trials to all double target green trials, regardless of whether double target green trials were performed correctly or not. While there was considerable variance, modulation indices across our sample averaged 0.09 ± 0.24, and were significantly distributed above zero (*t*-test, *t*(31) = 2.14, *p* < 0.05). Thus, on average, express visuomotor responses tended to be ∼10% smaller on double target green than double target red trials.

We then subdivided double target green trials based on whether monkeys correctly reached to the green stimulus or incorrectly reached to the red stimulus. There were 29 comparisons with sufficient numbers of both correct and incorrect double target green trials. The modulation index for comparison B compared double target red to correctly performed double target green trials; across our sample this modulation index averaged 0.17 ± 0.29 and was significantly distributed above zero (*t*-test Bonferroni-corrected for multiple comparisons, *t*(28) = 3.15, *p* < 0.005). On average, express visuomotor responses tended to be ∼20% smaller on correctly performed double target green than double target red trials. In contrast, comparison C compared double target red to incorrectly performed double target green trials; across our sample this modulation index averaged −0.06 ± 0.23 and was not significantly different from zero (Bonferroni-corrected *t*-test, *t*(28) = −1.4, *p* = 0.34). Finally, since comparisons B and C could be derived within the same session, we found that the modulation indices for comparison B tended to be significantly greater than for comparison C (Bonferroni-corrected paired *t*-test, *t*(28) = 4.22, *p* < 0.001). In other words, we observed a greater degree of contextual modulation on correct vs incorrect double target green trials. Overall, and as in humans, we found that the magnitude but not latency of the express visuomotor response changed with the context of the task (Gu et al., 2016; Contemori et al., 2021b, 2022a). Furthermore, as with anti-reaches (Gu et al., 2016), larger express visuomotor responses preceded incorrect responses.

## Discussion

We focused on the initial recruitment of upper limb muscles in monkeys reaching towards emerging targets, using a paradigm which readily elicits express visuomotor responses in humans (Kozak et al., 2020). Express visuomotor responses in monkeys exhibited the response properties known in humans: its timing is more time-locked to target emergence than movement onset, and its magnitude is both inversely related to RT and liable to dampening via contextual control. Our results establish how rapidly the visual-to-motor transformation is completed in the rhesus macaque brain, place constraints on the timing of premotor events, and set the stage for neurophysiological experiments that can directly compare the characteristics of such premotor events to those seen in the motor periphery.

### Comparisons of express responses in humans and monkeys

Rhesus macaques offer an excellent animal model for understanding rapid visual-to-motor transformations, due to the many homologies in foveate vision, implicated brain networks, and the musculoskeletal anatomy of the eyes, neck, and upper limb. Humans and macaques generate a variety of expedited visuomotor responses in the lab, including express saccades (Fischer and Weber, 1993), microsaccades (Hafed and Ignashchenkova, 2013; Tian et al., 2016), express visuomotor responses in neck muscle (Goonetilleke et al., 2015), reaches to rapidly moving stimuli (Perfiliev et al., 2010), and ocular following responses (Miles et al., 1986; Gellman et al., 1990). A common finding is that such responses evolve ∼20-30 ms earlier in macaques than humans, which is broadly consistent with the 3/5th rule (Schroeder et al., 2004) for quicker sensory processing in macaques versus humans. Consistent with this, express visuomotor responses in macaques started at ∼65 ms, compared to 80-100 ms in humans.

A notable exception for earlier responses in macaques vs. humans is when subjects adjust reaching trajectories in mid-flight to a displaced visual target. Humans can initiate such on-line corrections within ∼150-200 ms (Gaveau et al., 2014), even without conscious perception (Goodale et al., 1986). The response latencies for on-line corrections in macaques are generally longer than in humans, ranging between 200-250 ms (Georgopoulos et al., 1983; Archambault et al., 2009, 2011; Pastor-Bernier et al., 2012; Dickey et al., 2013), although latencies comparable with humans have been reported (Ames et al., 2019). The generally slower responses in macaques likely arises due to an task-imposed emphasis of accuracy over speed, so that animals were not responding as quickly as possible. Work by Perfiliev and colleagues (2010) compared ‘interceptive’ reaching responses to rapidly moving targets of humans, monkeys, and cats, and clearly showed that all three species can initiate reaches in less than 150 ms, with limb muscle recruitment in humans starting at 90-100 ms. Such recruitment onsets are within the range of express visuomotor responses in humans, but Perfiliev and colleagues did not record EMG in cats or monkeys.

### Methodological considerations

A study by Lara and colleagues (Lara et al., 2018) recorded upper limb muscle activity in monkeys in a ‘quasi-automatic’ task, where monkeys intercepted a target moving radially on a touch screen. This task expedited RTs to ∼200 ms on average, which is similar to the reaction times we observed. As with the emerging target paradigm, the quasi-automatic task features a suddenly appearing moving visual target. Lara and colleagues recorded from multiple heads of deltoid in addition to other upper limb muscles, reporting the earliest changes in average muscle recruitment at ∼100 ms, with single-trial onsets as low as 90 ms. Although these response latencies are short, they are still ∼25-35 ms longer than those reported here. The failure for the quasi-automatic task to elicit express visuomotor responses may be due to a lack of precise temporal certainty about target presentation or the absence of implied motion, both of which are critical in the emerging target task (Kozak et al., 2020; Contemori et al., 2021a; Kozak and Corneil, 2021). The quasi-automatic task in the Lara study was also intermingled with other trial types that engender longer RTs, which may also lower the probability of express visuomotor responses. Finally, the Lara study employed eight potential target locations, whereas we employed two. We do not think this difference explains the absence of express visuomotor responses in the Lara study, as recent studies in humans have reported express visuomotor responses in paradigms with a similar or higher number of potential target locations (Gu et al., 2019; Kearsley et al., 2022; Selen et al., 2023) locations.

Caution is always required when comparing responses latencies across different laboratories. Stimulus luminance and spatio-temporal frequency impact the latencies of on-line reach corrections (Veerman et al., 2008; Kozak et al., 2019), the timing and magnitude of express muscle responses (Wood et al., 2015; Kozak et al., 2019; Kozak and Corneil, 2021), and on the timing and vigor of sensorimotor responses in the superior colliculus (SC) (Li and Basso, 2008; Marino et al., 2012; Chen et al., 2018). Determining whether limb muscle recruitment meets an “express” criterion should involve an assessment of trial-by-trial muscle recruitment over a range of RTs, lest averaged recruitment profiles be biased by trials with the shortest RTs. The trial-by-trial representations of EMG activity presented in the Lara study did not show consistently short recruitment latencies across trials, strengthening our contention that express visuomotor responses were not consistently elicited in their quasi-automatic task.

The emerging target paradigm provides a framework in which to study the most rapid visual-to-motor transformations for reaching, and our results show that the underlying phenomenon is distributed to many deltoid motor units in the macaque. The magnitudes of express responses compared to later waves of activity are more modest in monkeys than in humans. However, similarly-sized express visuomotor responses induce force transients or small movements toward the stimulus (Wood et al., 2015; Gu et al., 2016; Atsma et al., 2018). We speculate that the modest levels of recruitment may be because the deltoid muscles targeted here may not be prime movers when monkeys interact with a touchscreen. Future work recording from more muscles of the upper limb may find stronger express visuomotor responses, and experimental setups with a robotic manipulandum will allow loading of selected muscle, as done routinely in humans. We did observe small movements of the hand toward the earlier appearing red target on double target green trials, which we surmise arose from both the express visuomotor responses and subsequent periods of recruitment. The assessment of limb motion via the touchscreen is coarse, hence setups using robotic manipulanda or motor tracking will permit more precise assessment of the kinematic consequences of express visuomotor responses in monkeys. Future studies should also track eye movements. While human studies have shown that the eyes are stable at the time of stimulus presentation in the emerging target paradigm (Contemori et al., 2021a), what the eyes were doing at stimulus emergence in our study is not known. It is possible that the modest magnitude of express visuomotor responses could be due to saccades made around the time of target emergence, as such contemporaneous eye movements can decrease the vigor of visual responses in movement-related neurons in the superior colliculus (Chen and Hafed, 2017).

Double target green trials required the monkeys to suppress the tendency to look to the earlier-appearing red stimulus. We expected that RTs for such trials would be longer than the RTs for double target red trials, but somewhat surprisingly this was not the case if both correct and incorrect trials were pooled together. Future variants of this task may benefit from longer delays between the emergence of the red or green stimuli, and on-line identification of incorrect reaches. In our data, the incorrect reaches themselves were often quite small in amplitude, hence the animals could be rewarded on trials where they initially moved in the wrong direction so long as they eventually reached the target. That being said, arm movements have substantial inertia, and in such systems establishing the criteria used to distinguish “incorrect” from “correct” movements necessitate a degree of subjectivity. Determining such criteria is all the more pernicious given evidence that the express visuomotor response itself, whether on upper limb or neck muscles, imparts sufficient forces to subtly move the arm or head (Corneil et al., 2008; Gu et al., 2016). Regardless, reach trajectories (**Fig. 5C**) show that both animals rapidly corrected incorrect movements in mid-flight, which is consistent with them consolidating task instruction. The impact of task context on the express visuomotor response to the red stimulus was both modest and variable (**Fig. 6**), but the average magnitude resembled that seen on monkey neck muscles performing pro- or anti-saccades (Chapman and Corneil, 2011). Contextual modulation of the express visuomotor responses is likely something that can be optimized in future experiments, for example by employing various endogenous or exogenous cueing strategies that potentiate the phenomenon of express visuomotor responses in the human arm (Contemori et al., 2021b, 2022a) or monkey neck (Corneil et al., 2008).

### Implications of our results

Express visuomotor responses are thought to arise from a subcortical tecto-reticulo-spinal circuit that runs in parallel with corticospinal circuits (Boehnke and Munoz, 2008; Pruszynski et al., 2010; Corneil and Munoz, 2014; Contemori et al., 2022b). This hypothesis has never been directly tested in monkeys, as the lone reaching study inactivating the SC during a reaching task (Song et al., 2011) did not record limb EMG and used a delayed response task that would not engender express visuomotor responses. Considerable circumstantial evidence implicates the visual response within the intermediate and deep layers of the superior colliculus (dSC) as being key in driving express responses. The timing and vigor of the visual response, when summed with pre-existing preparatory activity, drives express saccades (Edelman and Keller, 1996; Dorris et al., 1997; Sparks et al., 2000; Dash et al., 2018). Visual responses in the dSC are also correlated with the magnitude of express visuomotor responses in the neck, leading such responses by an appropriate efferent lag (Rezvani and Corneil, 2008). Reach-related neurons in the dSC display short-latency visual responses (Werner et al., 1997a, 1997b) and correlate with upper limb muscle activity (Werner et al., 1997a; Stuphorn et al., 1999), although the properties of visual responses in reach-related neurons have not been systematically investigated. Our speculation of a subcortical route initiating visually-guided reaches to a punctate target is consistent with recent stimulation results in humans (Divakar et al., 2022), and parallels a growing appreciation for the role of subcortical pathways in manual following responses to whole-field motion (Saijo et al., 2005) or flashes (Nakajima et al., 2021).

While we acknowledge the shortcomings in comparing results across laboratories and experimental tasks, the express visuomotor responses reported here on arm muscles occur at an interval when neural activity in the primary motor cortex is thought to be in a null space that is not tied to motor output (Lara et al., 2018). Indeed, one of the main conclusions for the Lara study mentioned above is that a mandatory transition of neural activity within motor cortex from a null-space to a movement-potent subspace precedes motor output by ∼20-30 ms. Our results present two alternatives which can frame future experiments.

First, if our hypothesis is correct, a subcortical route through the dSC can drive express visuomotor responses when M1 is in a null space. An intriguing corollary to this alternative, given ascending connections from the SC to thalamic areas that themselves project to motor cortices (Benevento and Fallon, 1975; Matelli and Luppino, 1996), is that it is signaling from the SC to the motor cortex that drives the transition of cortical activity from null to movement potent subspaces (Ames et al., 2019), which presumably could allow cortical activity to drive subsequent muscle recruitment that is informed by the consequences of the preceding express visuomotor response. Ascending signals from the SC through the thalamus have been long established as being critical for relaying efference copy signals for eye movements (Sommer and Wurtz, 2002), and implicated in prioritizing spatial locations for target selection for reaching (Song et al., 2011) and for higher order processing (Lovejoy and Krauzlis, 2010; Bogadhi et al., 2021; Peysakhovich et al., 2023).

Alternatively, could the transition of neural activity from a null to movement potent subspace in M1 simply occur earlier in the emerging target paradigm? This line of reasoning surmises that signaling along a cortico-spinal or cortico-reticulo-spinal route drives express visuomotor responses. If so, visually-related activity would have to evolve in motor cortex ∼10-20 ms earlier than the express visuomotor responses we report, so at around 50 ms. There are reports of visual responses in the macaque primary motor cortex at such latencies (Murphy et al., 1982; Lamarre et al., 1983; Crammond and Kalaska, 1994; Riehle et al., 1997; Cisek and Kalaska, 2005; Ames et al., 2019), but comparisons to these reports are again hindered by the use of tasks or visual stimuli that in retrospect may not have been optimized to elicit express responses, and various filtering settings or analysis schemes that complicate timing comparisons. Examination of the rasters of single-unit data from such reports, when available, also do not always reveal the trial-by-trial consistency that we observed in the motor periphery. One intriguing exception comes from the work of Reimer and colleagues (Reimer and Hatsopoulos, 2010), who studied motor cortical activity while NHPs adjusted the trajectory of ongoing limb movements to jumped visual targets. When one accounts for an acknowledged timing error, motor cortex activity changed consistently ∼50 ms after the target jump. While this theoretically shows that signaling related to visual signal onset could arrive in time within motor cortices, we note that express visuomotor responses in humans evolve ∼10-15 ms earlier (and hence ∼80 ms or less) when evoked when the arm is already in motion (Kozak et al., 2019), presumably due to disengagement of a control policy for postural stability (Cluff and Scott, 2016).

## Acknowledgements

The authors thank Katherine Faubert, Rhonda Kersten, Kristy Gibbs, Darren Pitre, and Matthew Reeves for their technical assistance and animal husbandry support, and Kevin Barker for his apparatus design and production expertise. We thank Tim Carrol, Sam Contemori, Rechu Divakar, Sebastian Lehmann, and Gerald Loeb for feedback on earlier versions of this manuscript.

